# Threshold of somatic mosaicism disrupting the brain function

**DOI:** 10.1101/2023.12.30.573716

**Authors:** Jintae Kim, Sang Min Park, Hyun Yong Koh, Ara Ko, Hoon-Chul Kang, Won Seok Chang, Dong Seok Kim, Jeong Ho Lee

## Abstract

Somatic mosaicism in a fraction of brain cells causes neurodevelopmental disorders, including childhood intractable epilepsy. However, the threshold for somatic mosaicism leading to brain dysfunction is unknown. In this study, we induced various mosaic burdens in mice of focal cortical dysplasia type II (FCD II), featuring mTOR somatic mosaicism and spontaneous behavioral seizures. Mosaic burdens ranged from approximately 1,000 to 40,000 neurons expressing the mTOR mutant in the somatosensory (SSC) or medial prefrontal (PFC) cortex. Surprisingly, just ∼8,000-9,000 neurons expressing the MTOR mutant were sufficient to trigger epileptic seizures. Mutational burden correlated with seizure frequency and onset, with a higher tendency for electrographic inter-ictal spikes and beta- and gamma-frequency oscillations in FCD II mice exceeding the threshold. Moreover, mutation-negative FCD II patients in deep sequencing of their bulky brain tissues revealed somatic mosaicism of mTOR pathway genes as low as 0.07% in resected brain tissues through ultra-deep targeted sequencing (up to 20 million reads). Thus, our study suggests that extremely low levels of somatic mosaicism can contribute to brain dysfunction.

## Introduction

Somatic mosaicism (or mutations) which can arise during development or over aging, are increasingly recognized as an important genetic causes of brain disorders like focal epilepsy, autism, schizophrenia, and Alzheimer’s disease.^1^ While these conditions show widespread brain dysfunction, somatic mutations are found in a small fraction of brain cells such as neurons,^2,3^ astrocyte,^4^ oligodendrocyte,^5^ and microglia,^6^ even in specific brain areas. Although a single cell with harmful somatic mutations is unlikely to cause widespread brain dysfunction in non-neoplastic neurological diseases, the exact level of somatic mutation burden that disrupt the brain function remain unexplored yet.

Focal cortical dysplasia (FCD) is a focal malformation of cortical development and the most common cause of intractable childhood epilepsy subjected to epilepsy surgery.^7^ Among the various subtypes of FCD, FCD type II (FCDII) is a prominent one characterized by cortical malformation involving dyslamination and the presence of dysmorphic neurons^8^. This specific subtype of FCD is primarily attributed to somatic mosaicism, which leads to the abnormal activation of mTOR kinase within the dysmorphic neurons in the affected focal cortical region.^2^ We and other group have demonstrated that this somatic mosaicism often can be detected at levels as low as 1% of mutational burden (or variant allele frequency, VAF) in brain tissues extracted during surgical procedures.^3,9,10^

Notably, such a low level of somatic mosaicism is also adequate to induce epileptic seizures in a mouse model of FCDII, created through *in utero* electroporation of human mTOR mutant or CRISPR/Cas9 expressing vectors.^2,11,12^ However, it is essential to acknowledge the current limitations of sequencing technology that hinder the detection of low-level (e.g. less than 5%) pathogenic somatic mosaicism in patient’s tissues exists.^13^ Consequently, the precise threshold for somatic mutation that results in overall brain dysfunction remains uncertain yet.

## Materials and methods

### Patient ascertainment

This study included two patients with mutation negative FCDII diagnosis in deep gene panel sequencing,^14^ who had undergone epilepsy surgery at Severance Children’s Hospital, Yonsei University College of Medicine. Patients underwent comprehensive evaluation with video-EEG monitoring, high resolution MRI, fluorodeoxyglucose-positron emission tomography (PET), and subtraction ictal single-photon emission computed tomography (SPECT) co-registered to MRI to localize anatomic lesions. All patients met study entry criteria for FCDII.^15^ The pathologic diagnosis with FCDII was assessed by an experienced neuropathologist and reconfirmed for this study according to the consensus classification by the International League Against Epilepsy Diagnostic Methods Commision.^7^ This study was performed in accordance with protocols approved by Severance Hospital and the KAIST Institutional Review Board and Committee on Human Research, and all human tissues were obtained with informed consent.

### Mouse care and information

C57BL/6 mice (DBL, Korea) were used in experiments without sex discrimination, and they were housed in isolated cages with free access to food and water, and maintained in a room with a constant temperature of 23°C on a 12-h light-dark cycle with lights off at 7:00 p.m. All of the mice used in this study were healthy condition with normal immune status and had not involved in any previous test or drug treatment. All mouse experiments were approved by and performed in accordance with the guidelines set by the Institutional Animal Care and Use Committee (IACUC) of the KAIST.

### Mouse modeling and in vivo phenotype analysis

Model was generated according to previous articles with minor modification.^2,11^ Timed pregnant mice (E14.5) were used for generating somatic mosaicism of MTOR variant to one side of the hemicortex. For behavioral seizure analysis, every mouse involved in this experiment were undergone video seizure monitoring for 10 consecutive weeks (postnatal day 21∼84) to detect seizure onset and frequency. For EEG analysis, every mouse involved in this experiment were undergone video-EEG monitoring in 8 to 9 weeks after born, and the data was acquired and sampled for studying electrographic abnormalities including interictal spike and EEG spectra density. Detailed information about *in utero* electroporation, video and EEG monitoring and analysis is in **Supplementary material**.

### Imaging and three dimensional reconstruction analysis

After *in vivo* monitoring, mouse brain was perfused, fixed and sliced to 50∼60 sections for the subtotal imaging of whole cortex. After image acquisition, MATLAB software AMaSiNe was used to reconstruct the three-dimensional brain, calculate the number of mutant cells and automatically annotate each mutant cell onto Allen’s brain atlas.^16^ Detailed information about cell counting and volume estimation procedures are described in **Supplementary material**.

### Ultra-deep amplicon sequencing analysis

Genomic DNA was extracted from flash-frozen brain sections. Site-specific PCR amplification was done using specific primers to validate the identified mutations. **(Supplementary table 1)**. After amplification, products were sequenced on a Novaseq sequencer (Illumina, USA) for achieving high depth. Fastq files were preprocessed with fastp^17^ aligned to the reference genome of hg19 or mm10 and called for expected point mutation or indel read count. Detailed information about sample acquisition and library preparation is also described in **Supplementary material**

### Statistical analysis

All values in the figures are presented as the mean ± SEM. Grouped results were analyzed with Student’s t test, ANOVA where appropriate with GraphPad Prism7 (GraphPad Software). Non-linear fitting was done with logistic curve fitting built-in function:

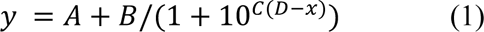

### Data availability

Data related with the findings of this study are available within this article and Supplementary material. Raw data are available upon reasonable request by author.

## Results

### Threshold number of mutation-carrying neurons triggering epileptic seizures in FCDII mouse model

To investigate the threshold of somatic mosaicism disrupting brain function, we employed an FCDII mouse model that exhibits mTOR somatic mosaicism in neurons through *in utero* electroporation of MTOR p.Leu2427Pro plasmid (**Fig 1A)**.^2^ These mice display spontaneous behavioral seizures during development.^2^ By adjusting the amount of injected plasmid within the range of 0.1 to 3μg, we were able to generate a spectrum of mosaic burdens in FCDII mice, ranging from approximately 1,000 to 40,000 neurons expressing the mTOR mutant, located in either the somatosensory (SSC) or medial prefrontal (PFC) cortex (**Supplementary Fig 1**).

**Figure 1.**
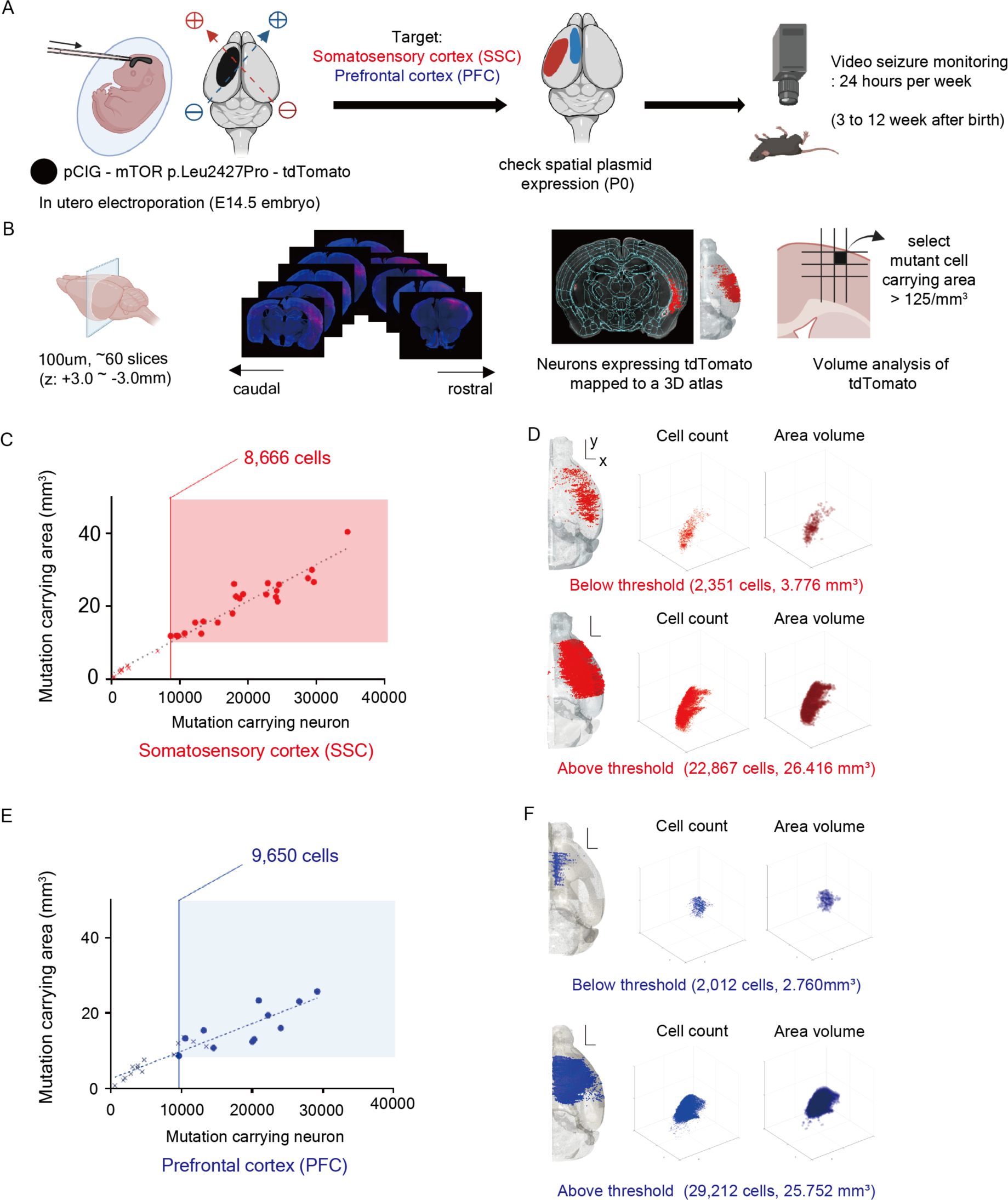
Threshold number of mutant neurons in the cortex triggering behavioral seizures. (**A**) Schematic figure of generating FCDII mouse model with mTOR somatic mosaicism, followed by video seizure monitoring. *In utero* (E14.5) electroporation of MTOR p.Leu2427Pro plasmid induced somatic mosaicism of mutant mTOR overexpression in mouse brain cortex with epileptic behavioral seizure. Onset and seizure frequency were video monitored. (**B**) Schematic figure of 3D reconstruction of mutant mTOR expressing cells using AMaSiNe. ∼60 coronal slices with 100um thickness were normalized to Allen’s brain atlas database and utilized for measuring cell numbers and volumes in reconstructed 3D brain. Reconstructed cell coordinates were further binned into 200um cube for describing mutant bearing region. (**C**) Mutant cells and the volume of carrying region was linearly correlated in somatosensory cortex. (Linear regression R=0.975, n=34. X ; no seizure, open circle : seizure) (**D**) Example of number and volume of mutant cells with below threshold (up) and above threshold (down) in somatosensory cortex. The axis follows Allen’s brain atlas data. (Scale bar x: 500um, y: 1mm) (**E**) Linear correlation found between mutant cells and the volume of carrying region in prefrontal cortex. (Linear regression. R=0.925, n=25. X ; no seizure, open circle : seizure) (**F**) Example of mutant cell carrying area in below threshold (up) and above threshold (down) in prefrontal cortex.

To quantify the number, location, and volumes of neurons expressing the mutant mTOR, we analyzed coronal slice images annotated to Allen’s brain atlas using AMaSiNe^16^ which reconstructs and calculates the neuron’s location within the brain’s standard coordinate framework (**Fig 1B**). Our analysis revealed a strong correlation between the number of mutant neurons and the volume of the area where these mutant neurons were distributed. Interestingly, behavioral seizures were only observed when there were more than 8,666 mutant neurons in the SSC, spanning an area of nearly 10 mm³ in the cortex (**Fig 1C and D**). Similarly, in the PFC, epilepsy development required at least 9,650 mutant neurons (**Fig 1E and F**). Considering that the isocortex in one hemisphere of the mouse consists of nearly 10 million cells,^18^ our findings suggest that a mere ∼0.1% of mutated neurons in the cortex are sufficient to trigger epileptic seizures, disrupting the overall brain function.

### Relationship between the level of mosaicism and the severity of epileptic seizures

Next, we investigated whether the number of mutant neurons above the threshold correlated with the seizure frequency and onset. We identified behavioral seizures by monitoring rearing, falling, and tonic-clonic seizures (Racine score 3-5) through 24-hour video surveillance from P21 to P90 (**Supplementary Fig 2**). Our findings revealed a non-linear sigmoidal relationship between seizure frequency and the number of mutant neurons in the somatosensory cortex (**Fig. 2A**). Furthermore, we observed an inverse correlation between seizure onset and the number of mutant neurons (**Fig. 2B**). A similar pattern was observed in the prefrontal cortex model, with a different plateau for average seizure frequency (**Fig. 2C and D**).

**Figure 2.**
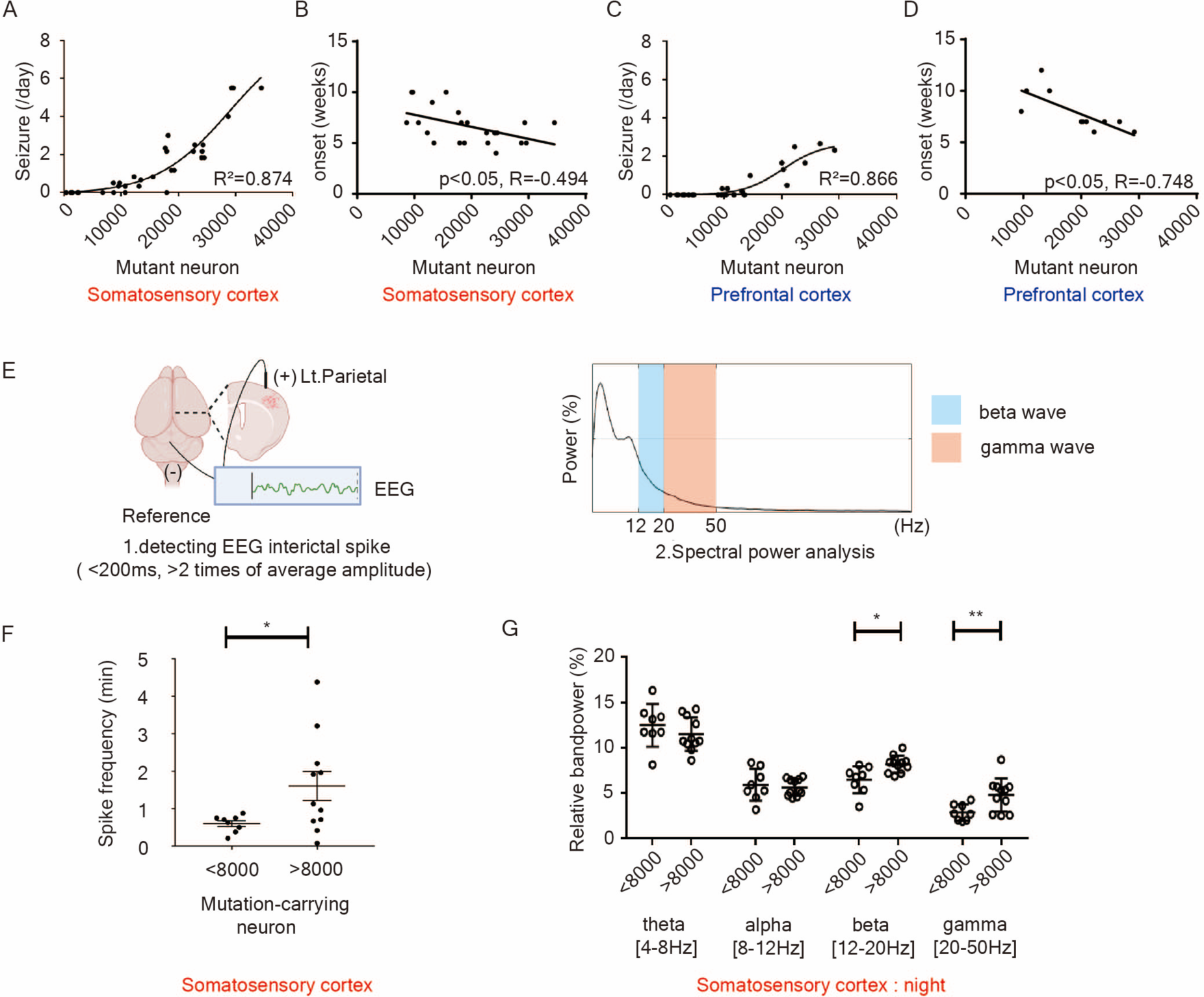
The relationship between the number of mutant neurons, behavioral seizure, and electrographic abnormality. (**A**) Non-linear sigmoidal relationship between mutant cell number and epileptic seizure frequency was found in somatosensory cortex model. (**B**) Negative linear relationship between mutant cell and epileptic seizure onset was found in somatosensory cortex model. (**C**) Relationship between cell number and seizure frequency in prefrontal cortex model. (**D**) Relationship between cell number and seizure onset in prefrontal cortex model. (**E**) Schematic figure of EEG monitoring in MTOR model (somatosensory cortex) (**F**) Difference of spike frequency between below (<8,000) and above (>8,000) threshold mutational burden (Unpaired Student t test with Welch’s correction, * *p* < 0.05, Low; n = 8, High; n=11) (**G**) Relative spectral density during night-time (24:00∼01:00) bandpower was calculated in 1-50Hz spectrum. Relative spectral density showed difference in beta (12∼20Hz) and gamma (20∼50Hz) spectrum during night. (Unpaired Student t test with Welch’s correction, * *p* <0.05, ** *p* <0.01)

For electrographic abnormalities, we conducted EEG analysis and found a higher interictal spike frequency in the SSC of FCDII mice with more than the threshold number of mutant neurons (**Fig. 2E and F**). High-frequency oscillations are known to contribute significantly to hyperactive neural field bursts.^19,20^ Consistent with this, we observed increased beta and gamma EEG spectral band power during the night (12 PM-1 AM) when the number of mutant neurons exceeded the threshold (**Fig. 2E and G, Supplementary Fig 3**). Taken together, our findings indicate that electrographic abnormalities begin to appear above the threshold, and the burden of somatic mosaicism positively correlates with the severity of epileptic seizures.

### Threshold of somatic mosaicism leading to epileptic seizures under physiological condition

Somatic mutations originating from neural stem cells affect both neurons and glial cells during brain development.^21^ Notably, prior research identified somatic mutations in MTOR in neurons and glial cells of FCDII patients.^3^ In FCDII model expressing mosaic mTOR mutant, *in utero* electroporation of mutant plasmids mainly led to the expression of mutant mTOR protein in neurons,^2^ which might underestimate the burden of somatic mosaicism actually presented in FCDII patients. To address this concern, we utilized another FCDII mouse model with somatic mosaicism of Tsc2, which leads to mosaic hyperactivation of mTOR kinase in both neural and glial lineages by CRISPR-Cas9 plasmid **(Fig 3A)**.^11^ Subsequently, we assessed the presence of behavioral seizures and measured the mutational burden of somatic mosaicism in the affected cortical regions. Given the expected low level of mutational burden in somatic mosaicism across different cell types, we performed ultra-high depth targeted amplicon sequencing with over 10 million read depth at the Tsc2 sgRNA target site to measure low-level mutational burden **(Fig 3B).**

**Figure 3.**
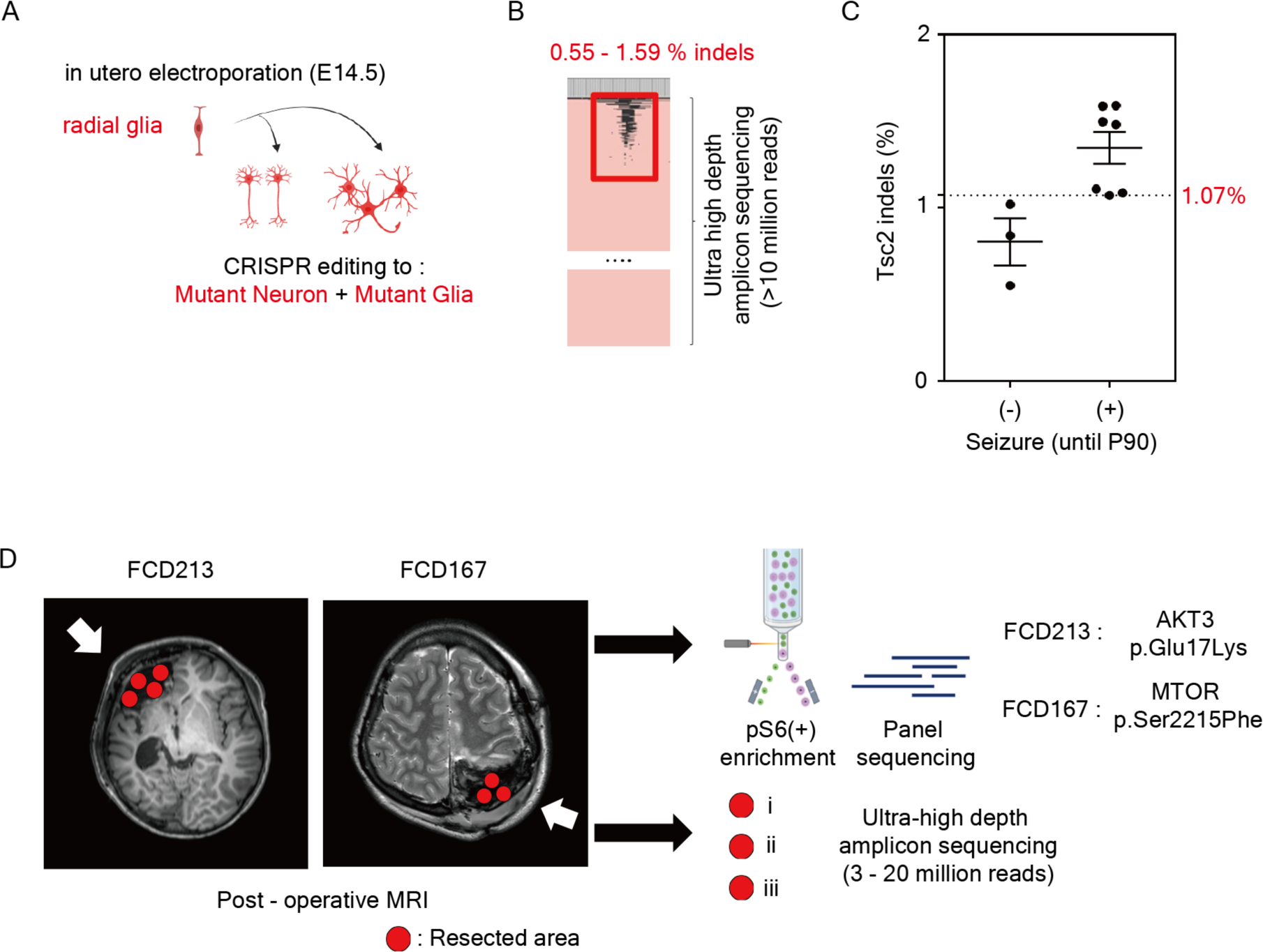
Extremely low level of somatic mosaicism under physiological condition. (**A**) Tsc2 CRISPR editing *in utero* electroporation result in somatic mutation of neuron and glial cells. (**B**) Multiclonal indels were observed in ultra-high depth amplicon sequencing data of bulky brain tissues carrying Tsc2 somatic mosaicism. (**C**) Percentage of sequencing reads with indels per total reads in Tsc2 mosaic mutation mice with or without seizures. Non-targeting indels were excluded. (**D**) Post-operative MRI of two FCD type II patients (FCD213 and FCD167), who were somatic mutation-negative for deep sequencing of their bulky brain tissues^14^. In both patients, somatic mutations in AKT3 or MTOR were detected as low level in deep sequencing of FACS enriched pS6(+) cells. In parallel, multiple samples of bulky brain tissues without FACS enrichment were applied for ultra-high depth amplicon sequencing with up to 20 million read depth.

As a result, we established FCDII models with Tsc2 somatic mutations, exhibiting various VAFs of indel reads ranging from 0.55% to 1.59%, corresponding to the same level of mutated cells in the resected brain cortex. Interestingly, behavioral seizures were only observed when the percentage of mutated cells in the resected cortex exceeded 1.07% **(Fig 3C)**. Therefore, considering the nature of brain somatic mosaicism, where somatic mutations arise from neural stem cells present in both neurons and glial cells during development, our findings suggest that as low as 1% of mutated cells can be sufficient to trigger epileptic seizures and brain dysfunction in FCDII patients.

### Presence of extremely low-level somatic mosaicism in mutation-negative FCDII

Finally, we investigated whether such a low level of somatic mosaicism is indeed present in FCDII patients who could not be genetically diagnosed through current deep-sequencing technology such as deep whole exome or gene panel sequencing with several hundreds to thousand read depths.

Despite recent updates in the classification and diagnostic criteria for FCDII,^8^ which now consider somatic mosaicism in mTOR pathway genes as a major genetic cause, approximately 40% of FCDII patients remain genetically unexplained.^3,9^ In cases where deep whole exome or gene panel sequencing did not identify somatic mosaicism in mTOR pathway genes in bulk brain tissues, we recently discovered that such mutations were detectable in Fluorescence-Activated Cell Sorting (FACS) enriched cells with mTOR hyperactivation and increased phosphorylated S6 (P-S6).^14^ To determine the precise mutational burden in these FCDII patients, we obtained multiple brain tissue samples from two FCDII individuals (FCD213 and FCD167) (**Fig 3D and Supplementary table 2**).

In these patients, enriched P-S6 cells exhibited the AKT3 c.49G>A (p.Glu17Lys) mutation in FCD213 and the MTOR c.6644C>T (p.Ser2215Phe) mutation in FCD167.^14^ We conducted ultra-high-depth amplicon sequencing with up to ∼19 million read depth in the resected bulk tissues and found VAFs of somatic mosaicism ranging from 0.07% to 0.60%, depending on the sampling site. The average VAF of somatic mutations was 0.25% in FCD213 and 0.50% in FCD167 in brain. Comparing with the matched blood sequencing data having less than 0.02% of false positive allele frequency(**Supplementary table 3**), our result are showing that confirmation of under 1% of variant allele frequency could be achieved using ultra high depth sequencing over million read depths.

Given that somatic mutations are typically present in one allele of somatic cells, these mutational burdens correspond to 0.5% of mutated cells in FCD213 and 1% in FCD167. Remarkably, following epilepsy surgery, both patients showed significant level of recovery, underscoring that even this extremely low level of mutation was epileptogenic in these cases. In conclusion, our results demonstrate that even less than 0.5% of somatic mosaicism burden is sufficient to cause widespread brain dysfunction in FCDII patients(**Table 1**).

**Table 1.** Ultra deep amplicon sequencing of mutation-negative FCDII patients and their surgical outcomes.

| **Surgical prognosis of FCD213 & FCD167** |  |  |  |
| --- | --- | --- | --- |
| **Patients** | **Seizure frequency(/day)** | **Seizure frequency(/day)** | **Seizure frequency(/day)** |
|  | **before surgery** | **6 months after surgery** | **3 years after surgery** |
| FCD213 | 26 | 0 | 0.03 |
| FCD167 | 20 | 0 | 0.01 |
| **FCD213 : AKT3 c.49G>A (p.E17K)** |  |  |  |
|  | **Total reads** | **Altered reads** | **Variant allele frequency (%)** |
| FCD213 – i) | 3,468,782 | 8,587 | 0.248 |
| FCD213 – ii) | 4,046,167 | 10,937 | 0.270 |
| FCD213 – iii) | 4,973,972 | 3,676 | 0.074 |
| FCD213 – iv) | 4,437,599 | 16,393 | 0.370 |
| **FCD167 : MTOR c.6644C>T (p.S2215F)** |  |  |  |
|  | **Total reads** | **Altered reads** | **Variant allele frequency (%)** |
| FCD167 – i) | 16,238,939 | 55,742 | 0.343 |
| FCD167 – ii) | 17,378,072 | 105,270 | 0.606 |
| FCD167 – iii) | 18,653,180 | 105,058 | 0.557 |

## Discussion

Here, we show that a specific threshold of somatic mosaicism leads to widespread brain dysfunction, including epileptic seizures in both mouse models and patients. In the FCDII mouse model, approximately 8,000-9,000 mutant neurons in the cortex, accounting for about 0.1% of mouse isocortical cells, are sufficient to disrupt normal brain function causing epileptic seizure and electrographic abnormalities. In the FCDII patients, even less than 0.5% of somatic mosaicism burden found in focal epileptogenic tissue is enough to cause widespread brain dysfunction including epileptic seizure In FCD II patients, the extent of brain lesion is known to be correlated with VAFs of somatic mosaicism.^10^ However, the relationship between mutational burden of somatic mosaicism and severity of epileptic seizure remains unclear. This study demonstrates that as the mutational burden increases, the epileptic seizure phenotype does not linearly correlate with mutational burden but follows a sigmoidal pattern, indicating a sudden state transition to daily epileptic seizures above a certain threshold. This may explain why clinical symptoms may not accurately reflect the level of variant allele frequency, as the severity plateaus within the detectable range of somatic mosaicism.

A somatic mosaicism burden of less than 0.5% is sufficient to trigger widespread brain dysfunction such as epileptic seizures in FCD II patients. One potential mechanistic explanation for this low-level threshold is that dysmorphic neurons expressing mTOR mutants in FCD II induce hyperexcitability in nearby non-mutant neurons through non-cell autonomous mechanisms, resulting in focal epileptic seizures.^22^ Nevertheless, further studies at the molecular level are needed to understand how this threshold level of mutant carrying neurons could produce widespread abnormal excitability to distant regions, eventually disrupting the whole brain. Moreover, FCD patients are showing additional neuropsychiatric symptoms including intellectual abnormalities and autistic symptoms.^3^ Further studies would also be needed to find how this few cells could disrupt the circuit level change in brain physiology.

Currently used sequencing technologies and analysis algorithms have limitations in detecting low-level somatic mosaicism, such as VAF less than 1%, due to sequencing errors, noises, and artifacts from library preparation.^23^ Although the threshold of somatic mosaicism dysregulating brain function can be variable depending on cell-types, regions, and disease phenotypes^9,24^, our study emphasizes the necessity of developing technology and tools to detect extremely low-level somatic mosaicism in various patients with different types of neurodevelopmental disorders. To address this, the world’s largest research initiative on somatic mosaicism, named ’Somatic Mosaicism across Human Tissues (SMaHT)^25^’ has recently been launched to accelerate the development of technology and tools for discovering somatic mosaicism including brain.

In summary, our study provides direct evidence of the extremely low-level threshold of somatic mosaicism leading to widespread brain dysfunction in both mice and humans.

## Acknowledgements

The authors thank the families that took part in this study and the neuropediatricians who referred the patients.

## Supplementary material

Provided in a separate file.

