## Supplementary Materials for "Threshold of somatic mosaicism disrupting the brain function"

**Supplementary Materials and methods (Page 2-5)**

**Supplementary Figures 1-2 (Page 6-7)**

**Supplementary Tables 1-4 (Page 8-13)**

### Supplementary Methods

#### *In utero* electroporation modelling of FCD type II

Timed pregnant mice (E14.5) were anesthetized using isoflurane (0.4 L/min of oxygen and isoflurane vaporizer gauge 3 during surgery operation). 2 μg/ml of Fast Green (F7252, Sigma, USA) combined with 0.1–3 mg of desired plasmids were injected to lateral ventricle of each embryo through uterus wall using pulled glass capillaries. Electroporation on embryo brain was done by discharging 40 V with the ECM830 electroporator (BTX-Harvard apparatus) in five electric pulses of 100ms with 900-ms intervals. For targeting somatosensory(SSC) cortex, positive electrode was placed onto plasmid inserted ventricle. Prefrontal cortex(PFC) was targeted by placing electrode onto contralateral ventricle, as plasmid would be transfected to midline area, which would develop to frontal cortex, anterior cingulate cortex, and anterior part of retrosplenial cortex. Successful regional targeting and plasmid expression was checked with fluorescence gun (Seahorse) and later confirmed by imaging analysis. pCAG-MTOR(p.Leu2427Pro)-IRES-tdTomato or sgRNA(Tsc2 exon2 target)-Cas9-IRES-Cre was used to develop MTOR or Tsc2 FCD model.

#### Epilepsy video monitoring analysis

In mouse seizure monitoring, mouse was video recorded for 12 hours, 2 times per week during 10 consecutive weeks (postnatal day 21~84). Mouse was located in a small open field mimicking the environment of cage, with providing water gel and food. Mouse showing seizure for more than 20 seconds with Racine score 3 to 5 was checked for seizure.

#### EEG surgery, monitoring and analysis

To detect EEG signals, epidural electrodes were inserted to somatic mosaicism model. In adult mice (over 7 weeks), mice were anesthetized using isoflurane (0.4 L/min of oxygen and isoflurane vaporizer gauge 3 during surgery operation). Skin above bregma was excised (radius 1cm) and small burr holes was introduced on their skulls without piercing the brain parenchyma. After then, electrodes were implanted into burr hole. The EEG probes were properly connected with screws and wires loaded onto these holes and fixed to the cranium with dental cement and resin. Electrode located on temporal lobes (AP–2.4 mm, ML±2.4 mm) were recorded using cerebellum as a reference. EEG signals were recorded for 24 hours at 8~9 weeks of age, at least 4 days after electrode implantation. Signals were amplified with RHD2000 amplifier chip recorded with RHD2000 USB interface board (Intan technologies).

Analysis was done with MATLAB customized code. For interictal spike, 1 minute samples for every 1 hour bin of total data (total 24 minutes) were selected for analysis, without severe digital noise. Spikes were defined with fast (<200 ms) epileptiform waveforms that were at least twice the amplitude of the background activity of total 1 hour bin. For spectral analysis, each 1 hour epoch during light and dark cycle (in 12AM and 12PM) were analyzed by Fourier transformation. Relative spectra were obtained by dividing power values of delta δ(1–4 Hz), theta θ(5–8 Hz), alpha α(9–12 Hz), beta β(12-20 Hz), or gamma γ(20–50 Hz) by total power values.

#### Fixation and cell counting analysis

Brains of mice were harvested after transcardiac perfusion with phosphate buffered saline and subsequent fixation using phosphate buffered 4% paraformaldehyde for approximately 12~24 hours. Fixed brain was divided into 50~60 coronal slices of 100um thickness (about 2.5~-3.5mm apart from bregma) using vibrating microtome (Leica VT1200). All sections were collected and placed on glass slides, mounted with prolonged gold antifade reagent with DAPI(P36931, Thermo Scientific). Images were acquired using a Zeiss slide scanner. Using MATLAB software AMaSiNe, they were aligned with anterior-to-posterior order and mutant cells with fluorescence were calculated for number and coordinate was automatically annotated onto Allen's brain atlas. Then, if there were two or more cells annotated in single voxel area ($0.008mm^3$), the voxel was included for volume estimation

#### Sample acquisition and library preparation for amplicon sequencing

For human sequencing study, samples acquired in previous study was used. To reflect the different part of pathologic FCD tissue, we acquired ~10mg of tissue from different areas of surgically resected brain, which was at least few centimeters apart from each other. For Tsc2 mouse sequencing study, mouse were undergone seizure monitoring for every week until as long as postnatal day 90. Every mouse were decapitated and brain was extracted. Hemispheric cortex with somatic mosaicism was resected for further study.

DNA from those samples were extracted using Qiamp Micro(<10mg) or Mini(>10mg) DNA kit according to manufacturer's protocol. Site-specific PCR amplification and library construction of multiple pathologic areas were done using region-specific primers for validating identified mutations in AKT3/MTOR(Human)/Tsc2(Mouse). Target sequences were PCR amplified with PrimeSTAR DNA Polymerase (Takara). 20 ng purified PCR product from the first amplification was annealed with barcode.

#### Statistical analysis

Details of statistical analysis used in this article is shown in **Supplementary Table 4.**

### Supplementary Figures

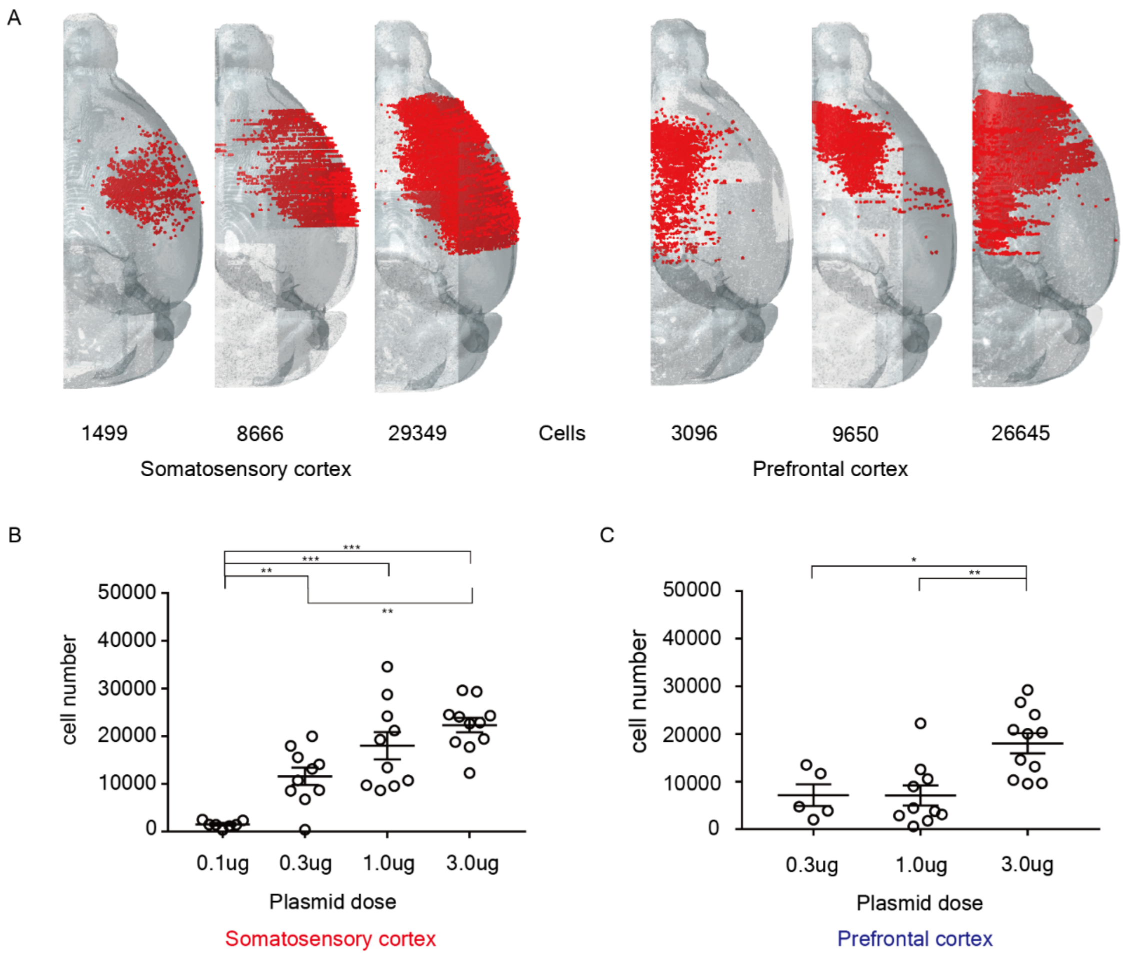


**Supplementary Figure 1. The number of mTOR mutant neurons is induced by in-utero electroporation of various amounts of plasmids.**

**a**, Examples of hemicortical distribution of mutant cell spectrum. The red dot represents mutant cell annotated on glass brain, using AMaSiNe code. **b-c**, Dose-dependent difference of mutant cell number is shown in some groups only. ANOVA with post-hoc t test. (*: p<0.05, **: p<0.01, ****: p<0.0001).

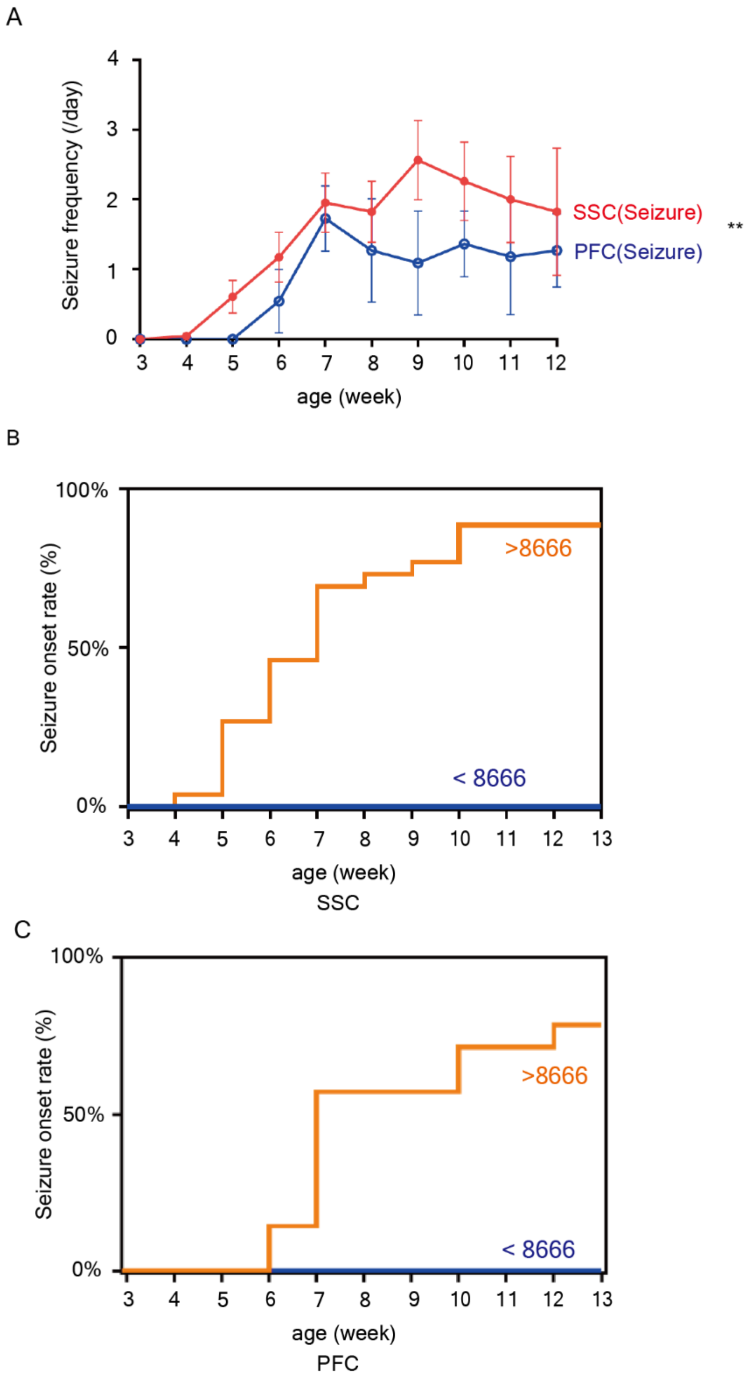


**Supplementary Figure 2. The seizure frequency and onset of FCDII mouse with mTOR mutant mosaicism.**

(**A**) Among FCDII models with seizures, site-specific weekly seizure frequency was statistically different. (Mean with S.E.M) (paired t-test) (**B-C**) with more than 8~9,000 cells mouse showed seizure during 24hr sampling period before $12^{th}$ week in 88.5% (left, somatosensory cortex) and 78.6% (right, prefrontal cortex) each. Epileptic seizure was not shown within mice having less than 8,666 cells until $12^{th}$ weeks after birth.

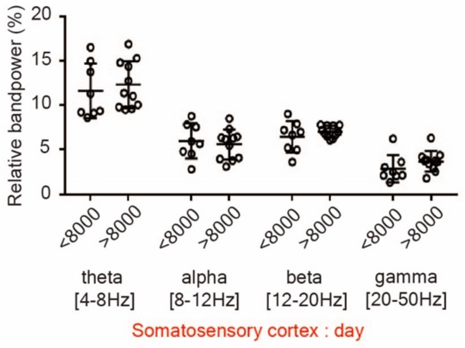


**Supplementary Figure 3. Spectral density analysis of EEG in FCDII mice during day**

Relative spectral density during day-time(12:00~13:00) bandpower was calculated in 1-50Hz spectrum (student t-test) showed non specific differences.

### Supplementary Tables

##### Supplementary Table 1 Primer for amplicon sequencing (AKT3, MTOR in human, Tsc2 in mouse)

| **Primer for Human** | **Forward(5')** | **Reverse(3')** |
|---|---|---|
| MTOR (S2215) | TGTTGCCATTTCAGGGTTTCT | GCAGCAACGGACATGAGTTT |
| AKT3 (E17) | CTTGCCACTGAAAAGTTGTTGAG | AACATGCGTGCTTTCCTCAT |
| **Primer for Mouse** | **Forward(5')** | **Reverse(3')** |
| Tsc2 (Exon 2) | AGACATTGGCCACTCACTCA | TTCTCTAGTACTGCCCTGCC |

Gene and exon specific primer set list for amplicon sequencing used in this article was presented.

**Supplementary Table 2. The clinical information of two patients with extremely low-level of somatic mosaicism.**

| Patient ID | Diagnosis | Sex | Age at surgery (years) | seizure onset | MRI/EEG | ASM used | Pathology |
|---|---|---|---|---|---|---|---|
| FCD 213 | FCD IIa | Male | 5.5 | 2 days | Cortical dysplasia involving right precentral and postcentral gyri | 4 | Cortical dyslamination, dysmorphic and giant neurons. |
| FCD 167 | FCD IIb | Female | 13.1 | 5 months | Diffuse cortical dysplasia in occipitoparietal lobe. extended to left precentral gyrus. | 3 | Cortical disorganization, dysmorphic neurons and balloon cells |

Both patients showed clinical seizures with early onset and severe symptoms. Radiological and pathological diagnosis correlated with typical FCD type II. ( FCDIIa: focal cortical dysplasia type IIa, FCDIIb: focal cortical dysplasia type IIb, EEG: electro-encephalography, ASM: anti seizure medication)

**Supplementary Table 3. High-depth amplicon sequencing of matched blood sample in this study (FCD213, FCD167)**

| Sample Name | Total reads count | Altered reads count | Variant allele frequency (%) |
|---|---|---|---|
| FCD213-Blood (AKT p.E17K) | 9,005,135 | 1,246 | 0.014 |
| FCD167-Blood (MTOR p.S2215F) | 22,393,820 | 2,803 | 0.013 |

In FCD patients, matched blood of each patient was also gone under amplicon sequencing as a matched control sample. In order to exclude the batch specific error, those were differently barcoded with brain samples but were sequenced in the same batch using Novaseq (Illumina), and also the library preparation was done under the same experiment along with PCR amplification.

**Supplementary Table 4. Detail of statistical analysis and related tools in this study (page 1)**

| Figure 1 C, 1 E (Linear regression) | | | | | |
|---|---|---|---|---|---|
| **Figure** | **Equation** | **R square** | **N** | **F** | ***p*** |
| Figure 1 C | $Y = 9.973*10^{-4}X+1.568$ | **0.950** | 34 | 608.3 | <0.0001 |
| Figure 1 E | $Y = 7.342*10^{-4}X+2.54$ | **0.856** | 25 | 136.9 | <0.0001 |

X = cell number, Y = volume ($mm^3$), N = sample number

| Figure 2 A, 2 C (Non-linear regression) | | | | |
|---|---|---|---|---|
| | **Sigmoid Function** | **R square** | **Linear Function** | **R square** |
| Figure 2 A | $Y=8.643/(1+10^{0.677*(2.895-X)})-0.088$ | **0.874** | $Y=1.481*10^{-4}*X-0.7763$ | 0.751 |
| Figure 2 C | $Y=2.698/(1+10^{1.251*(2.017-X)})-0.024$ | **0.866** | $Y=9.133*10^{-5}*X-0.4886$ | 0.758 |

X = cell number/10,000 , Y = seizure frequency (/day), sample numbers are same as 1C and 1E respectively.

| Figure 2 B, 2 D (Linear regression) | | | | | |
|---|---|---|---|---|---|
| **Figure** | **Equation** | **R square** | **N** | **F** | ***p*** |
| Figure 2 B | Y = -1.188*X+8.976 | **0.950** | 34 | 6.80 | <0.05 |
| Figure 2 D | Y = -2.224*X+12.18 | **0.560** | 25 | 11.44 | <0.01 |

X = cell number, Y = seizure onset (week)

| Figure 2 F (Unpaired T test) | | | | | | | | |
|---|---|---|---|---|---|---|---|---|
| **Figure** | **N (control)** | **Mean** | **SEM** | **N (Exp.)** | **Mean** | **SEM** | **df** | ***p*** |
| Figure 2 F | 8 | 0.601 | 0.079 | 11 | 1.605 | 0.392 | 10.8 | **<0.05 (0.029)** |

| Figure 2 G (Unpaired T test) | | | | | | | | |
|---|---|---|---|---|---|---|---|---|
| **Figure** | **N (control)** | **Mean** | **SEM** | **N (Exp.)** | **Mean** | **SEM** | **dF** | ***p*** |
| Figure 2 G Theta | 8 | 24.95 | 1.67 | 11 | 22.97 | 1.12 | 12.92 | 0.341 |
| Figure 2 G Alpha | 8 | 11.79 | 1.25 | 11 | 11.22 | 0.56 | 9.854 | 0.683 |
| Figure 2 G Beta | 8 | 12.91 | 1.07 | 11 | 16.33 | 0.56 | 10.72 | **<0.05 (0.017)** |
| Figure 2 G Gamma | 8 | 5.69 | 0.64 | 11 | 9.57 | 1.11 | 15.34 | **<0.01 (0.008)** |

| Figure 3 C (Mann Whitney test) | | | | | | | |
|---|---|---|---|---|---|---|---|
| **Figure** | **N (control)** | **Mean** | **SEM** | **N (Exp.)** | **Mean** | **SEM** | ***p*** |
| Fig 3 C | 3 | 0.268 | 0.040 | 7 | 0.414 | 0.022 | **<0.05 (0.017)** |

(Statistical details for supplementary figure is on the next page)

#### Supplementary Table 4. Detail of statistical analysis (page 2)

| Supplementary Figure 1 (ANOVA, post hoc t-test) | | | | | | | |
|---|---|---|---|---|---|---|---|
| **Supple1 B (dose)** | **N (0.1)** | **N (0.3)** | **N (1)** | **N (3)** | | | **ANOVA *p*** |
| | 7 | 7 | 10 | 11 | | | **<0.0001** |
| (post hoc) | 0.1 / 0.3 | 0.1 / 1 | 0.1 / 3 | 0.3 / 1 | 0.3 / 3 | 1/3 | |
| P value | **0.04** | **<0.0001** | **<0.0001** | **0.0496** | **0.0020** | 0.1106 | |
| **Supple1 C (dose)** | **N (0.3)** | **N (1)** | **N (3)** | | | | **ANOVA *p*** |
| | 5 | 9 | 11 | | | | **<0.01 (0.0014)** |
| (post hoc) | 0.3 / 1 | 0.3 / 3 | 1 / 3 | | | | |
| P value | 0.856 | **0.011** | **0.002** | | | | |

| Supplementary Figure 2 (Wilcoxon matched-pairs signed rank test) | |
|---|---|
| **Figure** | **P value** |
| Supple 2 A | **0.0039 (Two-tailed)** |

| Supplementary Figure 3 (Unpaired T test) | | | | | | | | |
|---|---|---|---|---|---|---|---|---|
| **Figure** | **N (control)** | **Mean** | **SEM** | **N (Exp.)** | **Mean** | **SEM** | **dF** | ***p*** |
| Supple 3 Theta | 8 | 23.23 | 2.17 | 11 | 24.65 | 1.59 | 13.73 | 0.607 |
| Supple 3 Alpha | 8 | 12.00 | 1.40 | 11 | 11.23 | 1.01 | 13.59 | 0.651 |
| Supple 3 Beta | 8 | 12.93 | 1.07 | 11 | 14.1 | 0.39 | 8.42 | 0.391 |
| Supple 3 Gamma | 8 | 5.78 | 1.08 | 11 | 7.42 | 0.69 | 15.34 | 0.226 |

**Software and tools used in this studies for statistical analysis**

| Name | Provider | Details |
|---|---|---|
| Prism 7.00 | GraphPad | Statistical tests |
| ZEN blue | Zeiss | Acquiring images from slide scanner (Zeiss) |
| MATLAB | Mathworks | Running AMaSiNe, Estimation of volume |
| AMaSiNe | VSNN lab, KAIST https://github.com/vsnnlab/AMaSiNe | Detection and Annotation of cell in 3D brain atlas |
| Biorender | https://www.biorender.com/ | Drawing illustration |
| Samtools | http://www.htslib.org/ | Amplicon sequencing analysis procedure |
| Burrows-Wheeler aligner | Broad institute | |
